# Expanding the range of editable targets in the wheat genome using the variants of the Cas12a and Cas9 nucleases

**DOI:** 10.1101/2020.12.09.418624

**Authors:** Wei Wang, Bin Tian, Qianli Pan, Yueying Chen, Fei He, Guihua Bai, Alina Akhunova, Harold N. Trick, Eduard Akhunov

## Abstract

The development of CRISPR-based editors having different protospacer adjacent motif (PAM) recognition specificities, or guide RNA length/structure requirements broadens the range of possible genome editing applications. Here, we evaluated the natural and engineered variants of Cas12a (FnCas12a from *Francisella novicida* and LbCas12a from *Lachnospiraceae bacterium*) and Cas9 for wheat genome editing efficiency and ability to induce heritable mutations in endogenous genes controlling important agronomic traits in wheat. Unlike FnCas12a, LbCas12a was able to induce mutations in the wheat genome in the current study, even though with a lower rate than that reported for SpCas9. The eight-fold improvement in the gene editing efficiency was achieved for LbCas12a by using the guide RNAs flanked by ribozymes and driven by the RNA polymerase II promoter from switchgrass. The efficiency of multiplexed genome editing (MGE) using LbCas12a was mostly similar to that obtained using the simplex RNA guides. A LbCas12a-MGE construct was successfully applied for generating heritable mutations in a gene controlling grain size and weight in wheat. We show that the range of editable loci in the wheat genome could be expanded by using the engineered variants of Cas12a (LbCas12a-RVR) and Cas9 (Cas9-NG and xCas9) that recognize the TATV and NG PAMs, respectively, with the Cas9-NG showing higher editing efficiency on the targets with atypical PAMs compared to xCas9. In conclusion, our study reports the set of validated natural and engineered variants of Cas12a and Cas9 editors for targeting loci in the wheat genome not amenable to Cas9-based modification.

## Introduction

Among considerations for designing genome editing experiments using an easily customizable CRISPR-Cas system, one of the important factors is the availability of target sequences with specific protospacer adjacent motifs (PAMs). The Cas9-based editor’s specificity is defined by the 20 nt-long target sequence located next to G-rich NGG PAM (Jinek et al., 2012), which limits availability of target sequences. The discovery of Cas12a nuclease (Zetsche et al., 2015) and engineered SpCas9 variants (Hu et al., 2018a; Nishimasu et al., 2018) allows to expand the number of editable target sites. The T-rich PAMs of Cas12a (TTTV or TTV) and the atypical NG PAMs of xCas9 and Cas9-NG allows for genome editing in the regions that might lack the G-rich PAMs needed for SpCas9. When the Cas12a nucleases were initially studied in human cells, contrary to the FnCas12a from *Francisella novicida* U112, only AsCas12a from *Acidaminococcus sp. BV3L6* and LbCas12a from *Lachnospiraceae bacterium ND2006* induces detectable mutations (Zetsche et al., 2015). Since the length of the PAM (TTTV) for the AsCas12a and LbCas12a nucleases limits the possible target spaces for editing in genomes, the development of engineered AsCas12a (carrying mutations S542R/K607R or S542R/K548V/N552R) and LbCas12a (carrying mutations G532R/K595R or G532R/K538V/Y542R) with altered PAMs (TYCV or TATV) further broadened further the utility of the Cas12a system for genome engineering applications (Gao et al., 2017).

The application of CRISPR-Cas12a system has been widely studied in different plant species including *Arabidopsis*, rice, soybean, and tobacco (Begemann et al., 2017; Endo et al., 2016; Hu et al., 2017; Kim et al., 2017; Tang et al., 2017; Xu et al., 2017; Yin et al., 2017). In rice, the efficiency of LbCas12a-mediated gene editing was comparable to that obtained using Cas9, while the AsCas12a-induced mutations were hardly detectable (Tang et al., 2017). The FnCas12a, which has shorter PAM (TTV), was proved to induce robust DNA cleavage in multiple species, including such model species as rice and tobacco (Begemann et al., 2017; Endo et al., 2016; Wang et al., 2017; Wang et al., 2018a; Zhong et al., 2018). In wheat, until now, only two target sites in the exogenous gene GUS (β-glucuronidase) were edited using LbCas12a (Liu et al., 2020), with one of the targets showing the genome editing efficiency lower than that of Cas9.

The efficiency of multiplex genome editing (MGE) is another important consideration when choosing the CRISPR-Cas system. To edit multiple targets in a genome using the CRISPR-Cas9 system, multiple guide RNAs (gRNAs) should either be expressed each from its own independent promoter or be expressed as a long tandem gRNA array with individual units separated by spacers, which should undergo endogenous or exogenous RNA processing to produce functional RNA guides (Wang et al., 2016; Xing et al., 2014). The ability of Cas12a to process a precursor crRNA array (Fonfara et al., 2016; Zetsche et al., 2017) permits using it without need to separate each guide by a spacer sequence. Additionally, the short length of crRNAs makes the CRISPR-Cas12a system an ideal MGE tool, which was successfully applied to editing the genome of different species, including rice (Wang et al., 2017).

To expand the range of editable targets, one of the most broadly used nucleases, SpCas9, was engineered to recognize the NG PAM instead of NGG. The two engineered variants, referred to as xCas9 and Cas9-NG, were initially tested in human/mammalian cells (Hu et al., 2018a; Nishimasu et al., 2018), and later in other species, including some crops (Ren et al., 2019; Zeng et al., 2019; Zhong et al., 2019). The xCas9 nuclease was substantially less effective than SpCas9 for editing the exogenous GUS gene in wheat (Liu et al., 2020), and less effective than Cas9-NG for editing the endogenous targets in rice (Zeng et al., 2019; Zhong et al., 2019). However, the application of both xCas9 and Cas9-NG for engineering the endogenous gene targets in wheat was not reported.

In this study, we investigated the nuclease activity of the natural and engineered variants of Cas12a and Cas9 towards the targets in the wheat genome with the canonical and non-canonical PAMs using the simplex and multiplex gene editing constructs. Using Cas12a we have produced a stable mutant with heritable mutations in a gene affecting grain size and weight in wheat. To improve the efficiency of Cas12a-based editing in wheat, we have evaluated constructs with humanized and plant codon-optimized variants of Cas12a expressing guide RNAs under the control of the PvUbip promoter from switchgrass, and assessed the effect of ribozyme-based guide RNA processing on the efficiency of genome editing. We have demonstrated the ability of the engineered Cas12a and Cas9 nucleases with the altered PAMs to induce mutations at targets inaccessible to Cas9, expanding further the range of editable loci in the wheat genome.

## Results

### Comparison of LbCas12a and FnCas12a genome editing efficiency in wheat

Because AsCas12a showed lower gene editing efficiency in both rice and tobacco compared to LbCas12a (Bernabe-Orts et al., 2019; Tang et al., 2017), in this study, we focused on evaluating the genome editing efficiency of LbCas12a and FnCas12a, which recognize the TTTV and TTV PAMs, respectively (Figure 1A). In total, 17 RNA guides were designed for LbCas12a to target eight genes: *TaAn-1, TaGASR7, TaGS3, TaGSE5, TaGW2, TaGW7, TaPDS*, and *TaSPL16* (Table S1). The efficiency of genome editing for 10 gene targets assessed in the wheat protoplast, after correcting for transformation efficiency, was higher than 1% in at least one biological replicate (Figure 1B, Table S2), with the highest editing efficiency reaching 21.6%. Most of the mutations induced by LbCas12a were deletions longer than 3 bp located at the 3’ end of protospacers distal from PAMs (Figure 1C).

**Figure 1.**
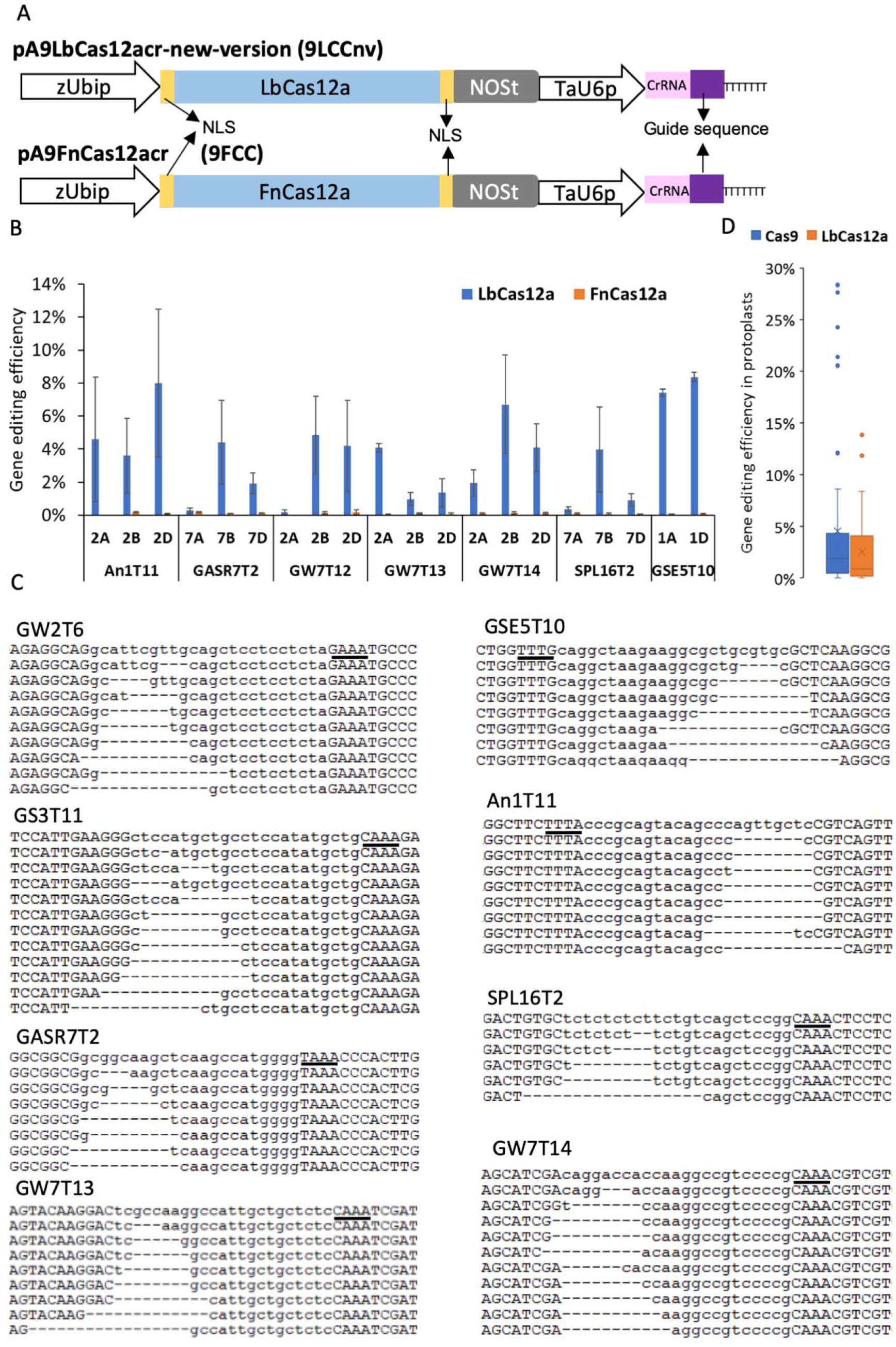
LbCas12a but not FnCas12a induces mutations in the wheat genome. **A)** A schematic illustration of plasmids pA9LbCas12aCr and pA9FnCas12aCr. Both the maize *Ubiquitin* promoter and wheat U6 promoter are shown with open arrows and marked as zUbip and TaU6p, respectively. The coding sequences of *LbCas12a* and *FnCas12a* are shown as blue rectangles. The nuclear localization signal peptide sequences that flank the *Cas12a* coding sequences are shown as yellow rectangles. The NOS poly A terminator is shown as grey rectangle and marked as NOSt. The direct repeat and guide sequence of crRNA are shown as pink and purple rectangle, respectively. The sequence of seven “T” bases is terminator for TaU6p. **B)** The comparison of gene editing efficiency between LbCas12a and FnCas12a. The gene editing efficiency was normalized by protoplast transformation efficiency. The bar plots showed the data as mean ± standard error. Each target had three biological repeats. **C)** The representative NGS reads generated for regions targeted by CRISPR-LbCas12a. The sequences of the wild type alleles are shown on the top. The PAM and target sequences are shown as underlined and lower-case letters, respectively. **D)** The comparison of mean gene editing efficiency for Cas9 (21 targets) and LbCas12a (12 targets) estimated for genes *TaGS3, TaGW7, TaPDS* and *TaGSE5* (Table S2 and S4). The duplicated target sites in the A, B, and D genomes were treated as independent targets. The gene editing efficiency of each target was normalized by protoplast transformation efficiency. The means of two or three biological replicates for each target were used to make the box and whisker plot.

Due to overlap of PAM sequences, the guides designed for LbCas12a should target the same genes when used with FnCas12a. However, using 16 guide sequences designed for LbCas12a and two guides specifically designed for the targets carrying the TTV PAM, we could not identify FnCas12a-induced mutations at the target sites in the wheat protoplasts (Figure 1B and Table S3).

To compare the gene editing efficiency between LbCas12a and Cas9, the CRISPR-Cas9 guides were designed for the coding sequences of the *TaGS3, TaGSE5*, and *TaPDS* genes. We also used the previously reported estimates of genome editing efficiency for the CRISPR-Cas9 guides targeting the *TaGW7* gene (Wang et al., 2019). The editing efficiencies for 21 Cas9 and 12 LbCas12a RNA guides were estimated to be 64% and 41% at the regions targeted by CRISPR-Cas9 and CRISPR-Cas12a, respectively, and both had mutation rates higher than 1% (Table S2 and S4). Overall, the gene editing efficiency of CRISPR-Cas9 was higher than that of LbCas12a in the wheat protoplasts (Figure 1D) The highest gene editing efficiencies observed for the LbCas12a targets within genes *TaGW7, TaGS3, TaGSE5*, and *TaPDS* were 6.7%, 13.8%, 8.4% and 3.74%, respectively. The highest gene editing efficiencies for the Cas9 targets within the *TaGW7, TaGS3, TaGSE5*, and *TaPDS* genes were 28.4%, 7.7%, 12.1% and 24.2%, respectively (Table S2 and S4).

### CRISPR-Cas12a-based multiplex gene editing

The MGE efficiency of the CRISPR-Cas12a system was investigated by transforming the wheat protoplasts using the constructs expressing LbCas12a and tandem arrays of three, four or eight crRNA units (Figure 2A and S2). The LbCas12a-induced mutations were detected for all targets included into the MGE constructs (Table S5). The editing efficiency of the multiplexed guides, except for GW2T6, from the LbCas12a MGE constructs, was comparable to that of the LbCas12a constructs carrying only one guide RNA (Figure 2B and S2, Table S5, *t*-test p-value > 0.05). The editing efficiency of the GW2T6 guide was significantly higher than those of the remaining guides and also increased with increase in the number of crRNA units in the tandem arrays of the LbCas12a MGE constructs (Figure 2B and S2).

**Figure 2.**
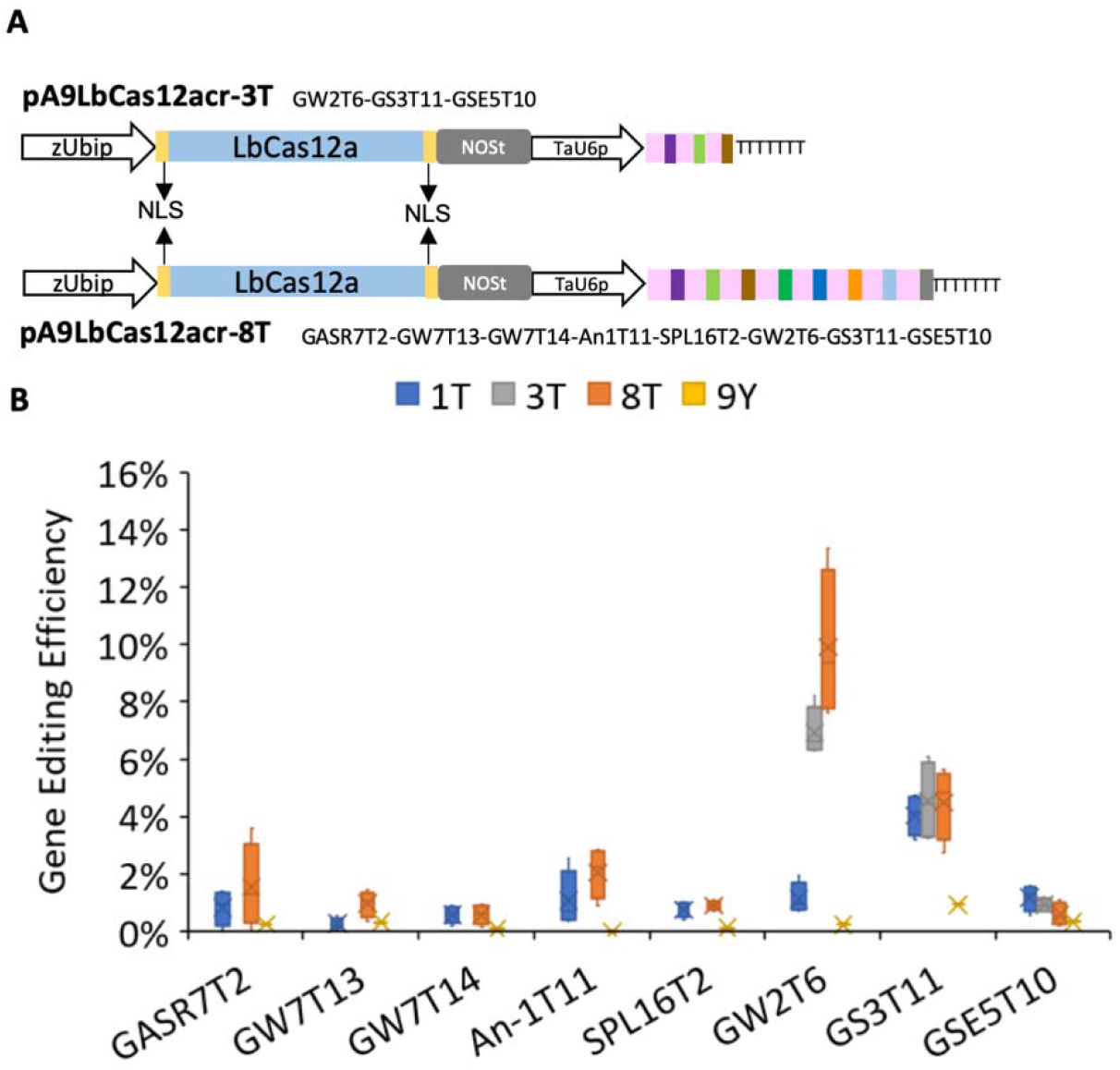
The gene editing efficiency of CRISPR-LbCas12a constructs with single and multiple gRNA units. **A)** Schematic illustration of the MGE CRISPR-LbCas12a constructs. The MGE constructs targeting three and eight targets are referred to as pA9LbCas12acr-3T and pA9LbCas12acr-8T, respectively. The order of guide sequences for different targets are shown next to the construct names. **B)** The box and whisker plots of gene editing efficiency for the CRISPR-LbCas12a constructs targeting one (1T), three (3T) and eight (8T) targets. The data generated from the protoplasts transformed with construct pA9eYFP was used as negative control (marked as 9Y). Because targets located in the A, B, and D genomes showed similar LbCas12a-induced mutation rates, data generated for all three genomes was pooled together for calculating the gene editing efficiency. The estimates for each target site were based on four biological replicates.

### A CRISPR-Cas12a-induced mutation in the *TaGW7-B1* gene changed grain shape and weight

To further validate the CRISPR-Cas12a system in wheat plants, we created 35 and 51 transgenic plants carrying the 9LCCnv-GS3T11 and 9LCCnv-8T constructs, respectively. All regions targeted by RNA guides in these LbCas12a-positive plants were genotyped by NGS. No mutations were detected in the 35 plants carrying the 9LCCnv-GS3T11 construct. Among 51 LbCas12a positive plants transformed with the 9LCCnv-8T construct, two plants (5384-1 and C3137-1) had mutations in the target site GW7T14, with no mutations detected in the remaining target sites. The T_0_ plant 5384-1 was heterozygous for 1-bp deletion in *TaGW7-B1*, and the T_0_ plant C3137-1 was heterozygous for 3-bp deletion in *TaGW7-D1* (Figure 3A). This was consistent with the segregation ratio of the mutated alleles in the T_1_ progeny of 5384-1 and C3137-1 plants (Figure 3B).

**Figure 3.**
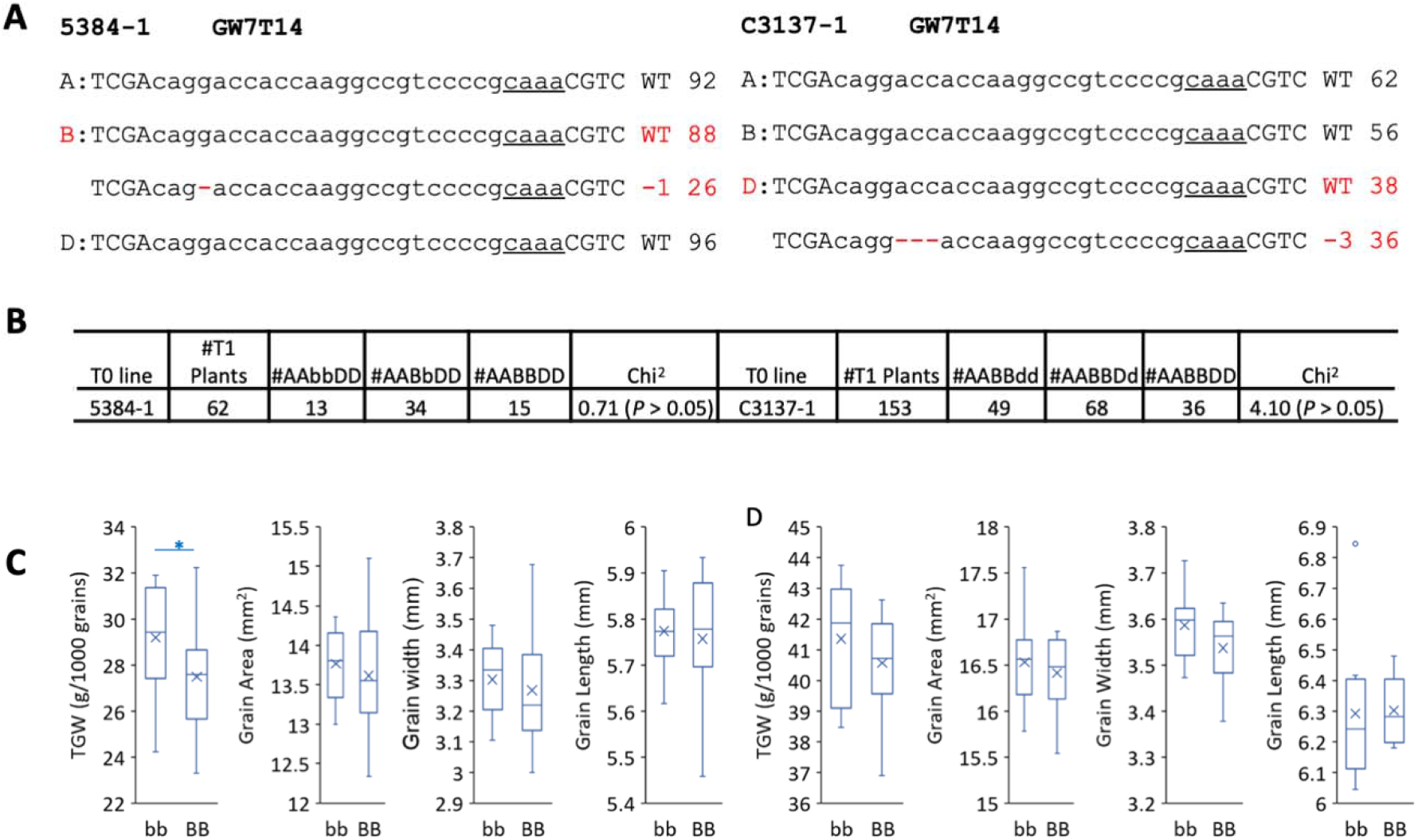
The phenotypic effects of heritable mutations induced by the *Cas12a* MGE construct. **A)** The aligned NGS reads flanking the target site GW7T14 and frequencies of wild-type and mutated reads in the T0 lines 5384-1 and C3137-1. WT stands for wild-type alleles in cv. Bobwhite; “-” sign and numbers after them represent number of nucleotides deleted. The PAM sequences are underlined. The deleted nucleotides are shown with red dashed lines. **B)** The segregation of Cas12a-induced mutated alleles in the *TaGW7* gene in T_1_ generation. The capital letters A, B and D and the lower-case letters a, b and d represent the wild-type and mutated alleles of *TaGW7* in the A, B and D genomes, respectively. The *χ*^2^-test for the expected Mendelian segregation ratio. **C)** and **D)** Box and whisker plots show the trait distribution for TGW, grain area, grain width, and grain length in T_1_ (**C**) and T_2_ (**D**) progeny of line 5384-1. The mean value of each group is shown as a “x” sign within the box-plot. Only the B genome genotypes with the *Cas12a*-induced mutations are shown. The capital letters and the lower-case letter represent the wild-type and mutated alleles, respectively. The T_1_ plants were derived from T_0_ line 5384-1, with 13 plants having genotype *AAbbDD*, and 15 plants having genotype *AABBDD*. All T_2_ plants were derived from T_1_ plant 5384-1-3 with 10 plants having genotype *AAbbDD*, and 11 plants having genotype *AABBDD*.

The effects of CRISPR-LbCas12a-induced mutations in the *TaGW7-B1* gene on grain size and thousand grain weight (TGW) were assessed in the T_1_ and T_2_ generation plants derived from 5384-1. The C3137-1 was excluded from further analyses because the 3-bp deletion does not cause frameshift in the coding sequence. In T_1_ generation, the TGW of *TaGW7-B1* homozygous mutants was increased by 6.2% compared to the wild-type lines segregated from the same T_0_ plants (*t*-test, *P* < 0.05) (Figure 3C). This result was confirmed by the 2% increase of TGW in the *TaGW7-B1* homozygous mutants compared to the wild-type plants in the T_2_ generation, though difference was not statistically significant (Figure 3D). While the grain length of *TaGW7-B1* homozygous mutants was not increased compared to the wild-type plants, the grain width was increased by 1.1% and 1.4% in the T_1_ and T_2_ generation, respectively. This increase of grain width was accompanied by the slight increase of grain area in the T_1_ and T_2_ generation plants. Though these increases in grain size were not statistically significant in both the T_1_ and T_2_ generation plants, the direction of phenotypic change in the mutant lines from both populations was the same, and also was consistent with the changes of TGW and grain size in the *TaGW7-B1* mutants previously created by our group using the CRISPR-Cas9-based genome editing (Wang et al., 2019).

### Improving the CRISPR-Cas12a-based genome editing efficiency in wheat

The editing efficiency of the CRISPR-LbCas12a construct for most genes was significantly lower than that of the CRISPR-Cas9-based constructs (Figs. 1D and 2B). In rice, it was shown that the LbCas12a’s editing efficiency could be significantly improved by using the maize ubiquitin promoter to drive expression of a crRNA flanked by ribozymes (Tang et al., 2017). To improve the CRISPR-LbCas12a gene editing capacity in wheat, we created a construct (hereafter referred to as pA9LbCas12acr-enhanced or 9LCCe), where the expression of crRNA flanked by two ribozymes was driven by the switch grass ubiquitin promoter (PvUbip) (Fig. 4A). Considering the ability of Cas12a to process CRISPR array into mature crRNAs, we tested a construct (henceforth pA9LbCas12acr-enhanced2 or 9LCCe2), in which the flanking ribozymes were removed and one extra direct crRNA repeat was added to the 3’ end of the crRNA protospacer (Figure 4A). In addition, the human codon optimized LbCas12a in 9LCCe and 9LCCe2 was replaced by the plant codon optimized LbCas12a (Tang et al., 2017) to create constructs henceforth referred to as 9LCCop and 9LCCop2, respectively (Figure 4A).

**Figure 4.**
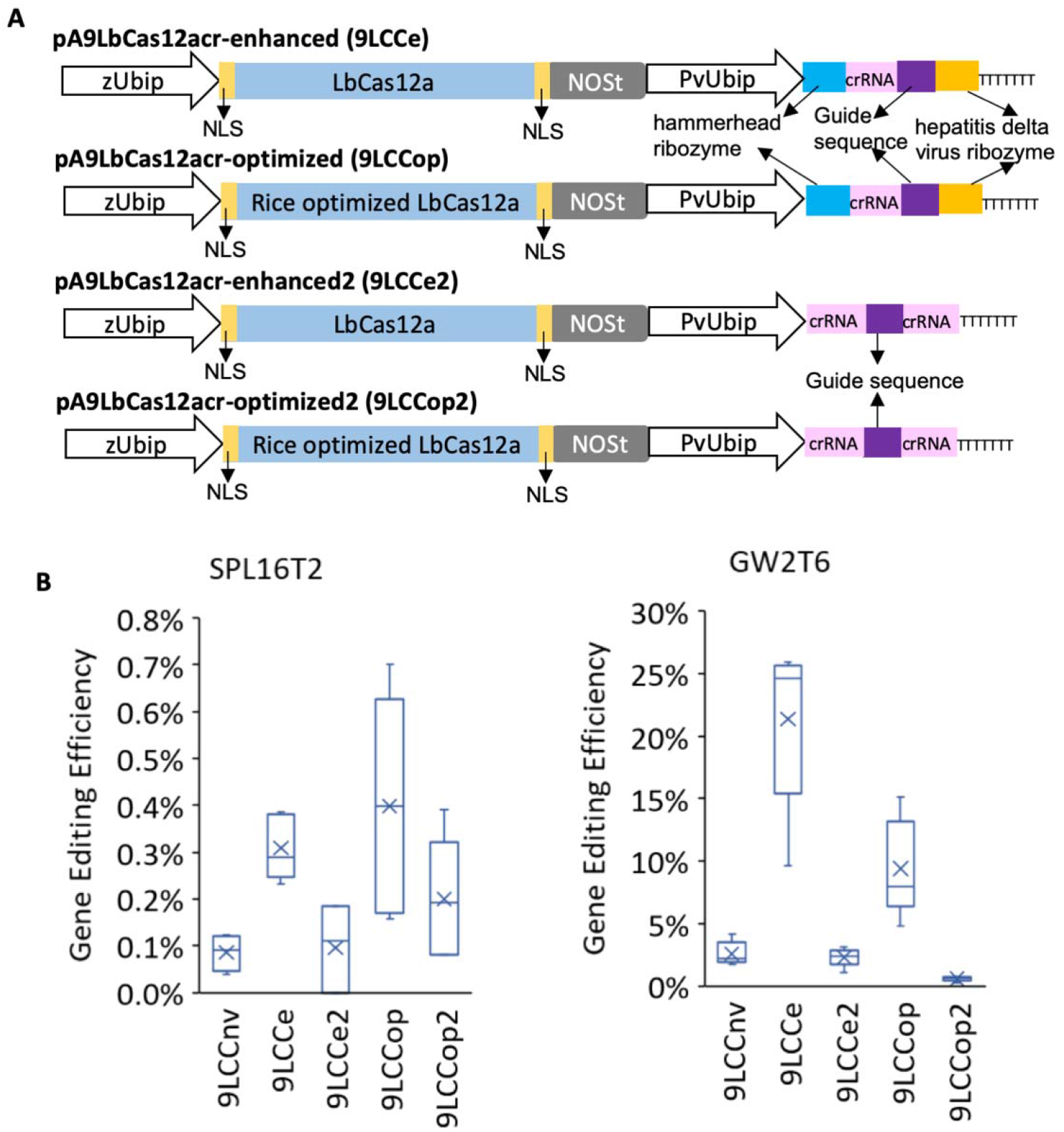
Improving the gene editing efficiency of LbCas12a. **A)** Schematic illustration of the modified LbCas12a constructs. Compared to 9LCCnv, all the constructs have the wheat U6 promoter replaced by the PvUbip promoter. 9LCCe and 9LCCop have hammerhead and hepatitis delta virus ribozymes flanking the 5’ and 3’ ends of crRNA to facilitate its processing; 9Lcce2 and 9LCCop2 have one extra direct repeat after the 3’ end of the guide sequence; 9LCCe and 9LCCe2 have the human optimized LbCas12a while 9LCCop and 9LCCop2 have the plant codon optimized LbCas12a. **B)** The comparison of gene editing efficiency among different versions of the LbCas12a constructs. The efficiencies of gene editing were estimated for the SPL16T2 and GW2T6 target sites conserved across all three wheat genomes. The estimates were based on four or five biological replicates. The gene editing efficiency was normalized by the protoplast transformation efficiency.

The CRISPR-LbCas12a crRNAs SPL16T2 and GW2T6 were subcloned into the modified constructs (Fig. 4A). Compared to the guides driven by the TaU6 promoter (9LCCnv construct), the gene editing efficiency of the SPL16T2 and GW2T6 crRNA flanked by ribozymes driven by the PvUbip promoter was improved by 3- and 8-fold, respectively (Figure 4B, Table S6). However, guides flanked by the crRNA scaffolds and driven by the PvUbip promoter did not show improvement in gene editing efficiency (Figure 4B, Table S6), indicating that the double ribozyme system improves the gene editing capacity of CRISPR-LbCas12a, likely by improving the efficiency of crRNA processing. The plant codon optimized Cas12a slightly improved the gene editing efficiency at target SPL16T2, but no improvement was observed at target GW2T6 (Figure 4B, Table S6), which indicates that codon optimization did not substantially affect the ability of CRISPR-LbCas12a to induce double strand breaks.

### The LbCas12a variant with the altered PAM induces mutations in the wheat genome

To broaden the editing capability of LbCas12a, we created a variant carrying mutations G532R, K538V, and Y542R (henceforth LbCas12a-RVR) that could recognize targets sites with the TATV PAMs (Figure 5A). As expected, the genome editing using LbCas12a-RVR at three target sites, GSE5T9, GW2T6 and PDST16, each having the TTTV PAM resulted in low mutation rate (Table S7). On contrary, by using LbCas12a-RVR in combination with two guides targeting sites with the TATV PAMs in the *TaAn-1* gene (Table S1), we detected mutations at both sites (Figure 5B) with the highest gene editing efficiency reaching 3.1% (Table S7).

**Figure 5.**
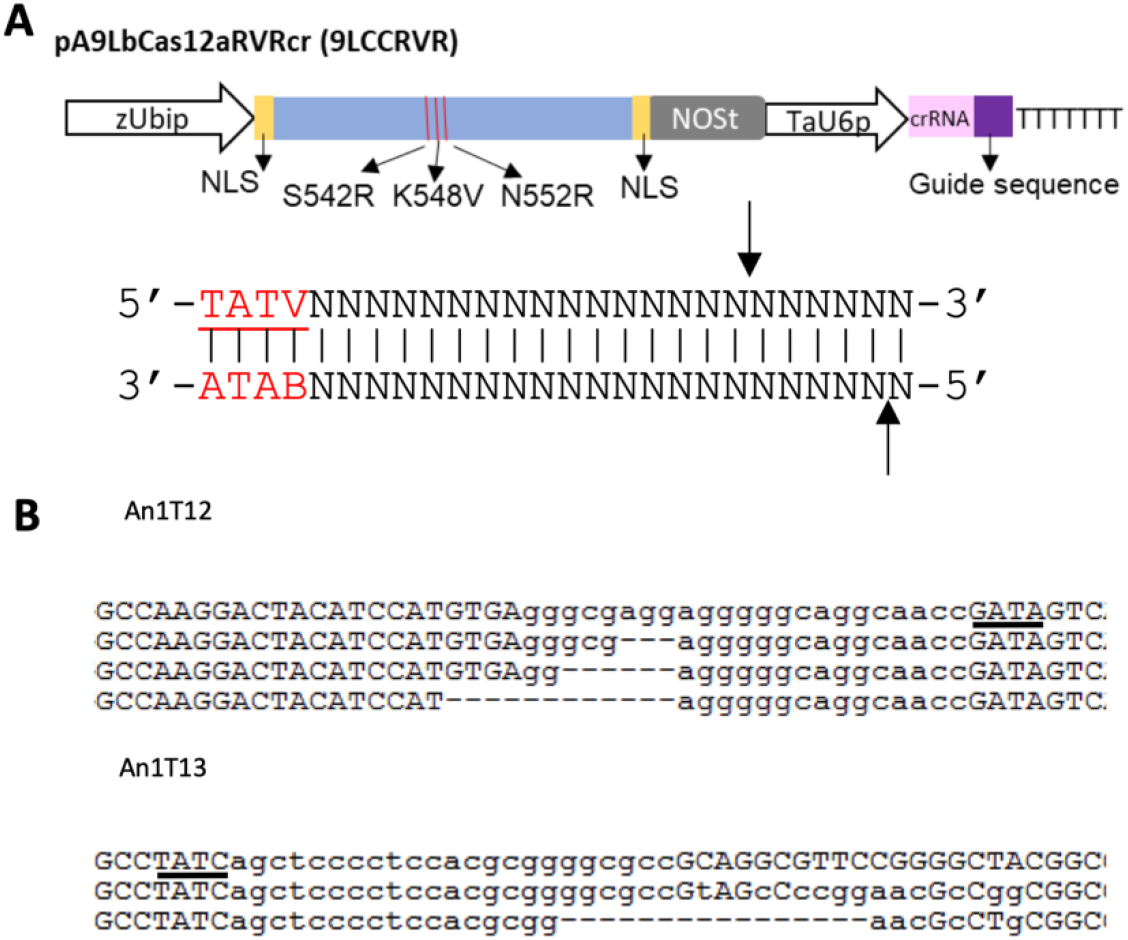
Engineered LbCas12a induces mutations in the wheat genome. **A)** Schematic illustration of plasmid pA9 LbCas12aRVRCr (9LCCRVR). The modified LbCas12a-RVR contains three amino acid substitutions, S542R, K548V, and N552R, which change PAM recognition specificity to “TATV”. **B)** The representative mutated reads of two targets on *TaAn-1* induced by CRISPR-LbCas12a-RVR. The first row for each target is wild type. The PAM sequence is underlined. The target sequences are shown in lower-case letters in the wild type sequence; the SNPs in the mutated reads of the An1T13 site are also shown as lower-case letters. The deleted nucleotides are shown as “-”.

### Off-target gene editing activity of LbCas12a and LbCas12a-RVR

It has been reported that both AsCas12a and LbCas12a show lower off-target activity in human cells compared to Cas9 (Kim et al., 2016a; Kleinstiver et al., 2016), likely due to the long PAM sequence. Here, we identified the possible off-target sites for the LbCas12a by comparing the target sequences with the reference genome IWGSC RefSeq v1.0 (The International Wheat Genome Sequencing Consortium (IWGSC) 2018). Among the analyzed 10 targets for LbCas12a and two targets for LbCas12a-RVR, only six targets had matching PAMs (Figure S3). Most of these possible off-target sequences were highly divergent within the 8∼10 bp from the PAM-distal end, and one of the possible off-target regions had SNPs located three and ten base-pairs after PAM (Figure S3). These mutations are expected to prevent LbCas12a from inducing mutations at the off-target sites. By sequencing these possible off-target sites, we demonstrated the lack of off-target editing activity for both CRISPR-LbCas12a or CRISPR-LbCas12a-RVR constructs (Table S8).

### Cas9-NG and xCas9 expand the range of genome editing targets in wheat

To investigate whether the xCas9 and Cas9-NG enzymes engineered to recognize NG PAMs could edit genes in the wheat genome, the maize codon-optimized Cas9 in construct pBUN421 was replaced by the synthesized maize codon-optimized xCas9 and Cas9-NG, henceforth referred to as pBUN421x and pBUN421NG, respectively. The gene editing ability of xCas9 and Cas9-NG was investigated by transforming them into the wheat protoplasts isolated from a wheat line constitutively expressing green fluorescent protein (GFP). Seven targets with the “NGN” PAMs were designed for the GFP coding sequence (Table S1). While both xCas9 and Cas9-NG induced mutations in the targets followed by NGG PAMs, they showed preference for different targets. Compared to xCas9, Cas9-NG showed eight times higher editing efficiency for target GFPT4 (Figure 6A). In contrast, xCas9 showed four times higher editing efficiency than Cas9-NG for target GFPT5 (Figure 6A). When compared to wild type Cas9, both xCas9 and Cas9-NG had lower editing efficiency for target GFPT13, which was followed by the NGG PAM (Figure 6A). For all four targets with the NGH PAMs, xCas9 showed lower editing efficiency than Cas9-NG (Figure 6A). The highest editing efficiency of 5.8% and 26% was observed for xCas9 and Cas9-NG on target site GFPT9 (Figure 6A and Table S9). While Cas9-NG induced mutations in targets GFPT7 and GFPT10 with the editing efficiency of 13.2% and 4.6%, respectively, no mutations on these targets were detected in the cells transformed with the xCas9 construct (Figure 6A and Table S9).

**Figure 6.**
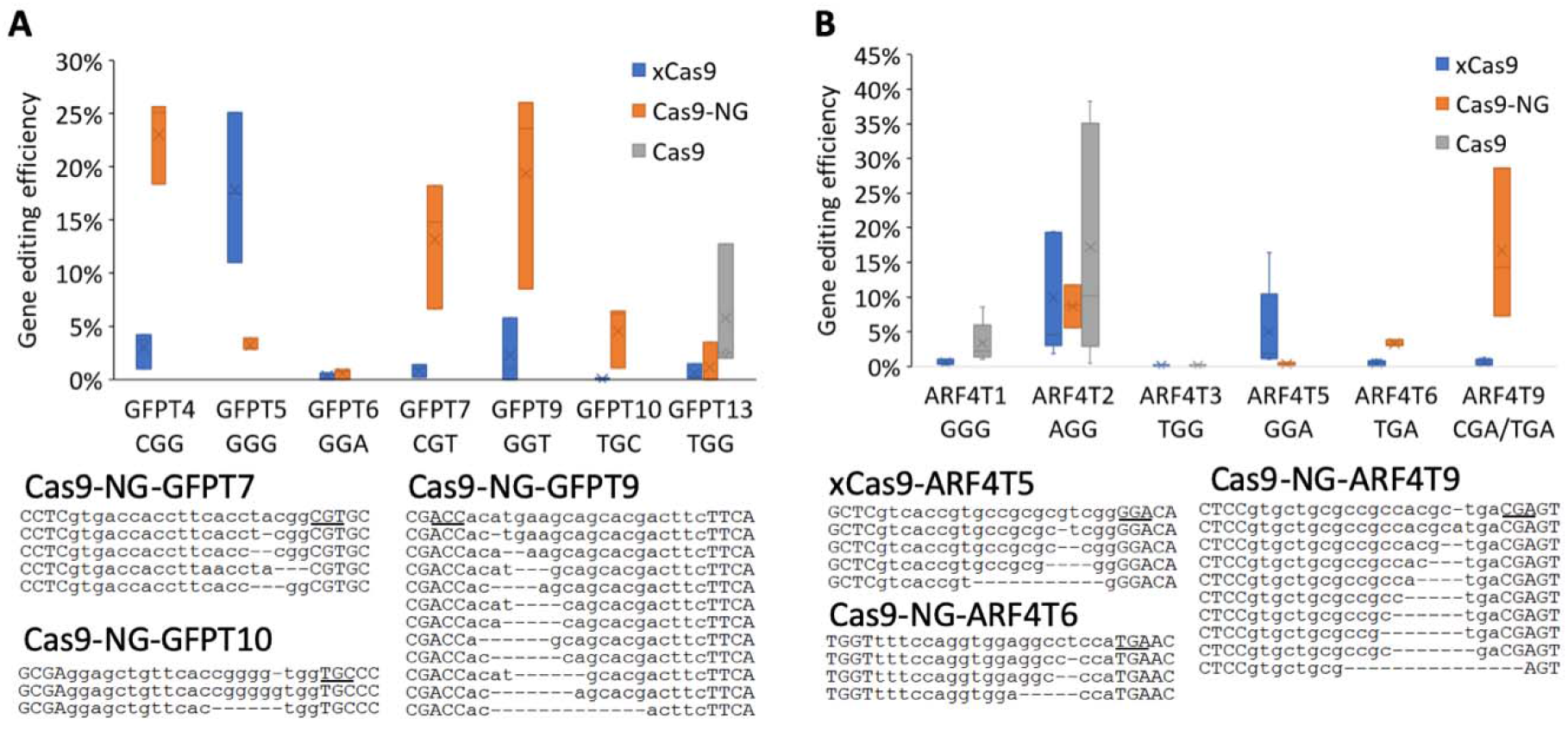
Mutations induced by the Cas9 variants in the wheat genome. **A)** Comparison of the xCas9 and Cas9-NG editing efficiency for targets located within the *GFP* gene. The estimates of gene editing efficiency are based on three biological replicates. **B)** Comparison of the xCas9 and Cas9-NG editing efficiency for targets located within the *TaARF4* genes. Because target sites were conserved across all three wheat genomes, the estimates of gene editing efficiency are based on reads pooled from all three wheat genomes. The gene editing efficiency was normalized by the protoplast transforming efficiency. The representative mutated reads for the Cas9-NG’s targets GFPT7, GFPT9, GFPT10, ARF4T6, and ARF4T9, and the xCas9’s target ARFRT5 are shown under the box and whisker plots. The first row for each target is wild type. The PAM sequence is underlined. The target sequences are shown in lower-case letters. The deleted nucleotides are shown as “-”.

To evaluate the ability of xCas9 and Cas9-NG to edit genes in the wheat genome, we designed guides for the *TaARF4* gene to target three regions with the NGG PAMs and three regions with the NGA PAMs (Table S1). Similar to target region GFPT13, both xCas9 and Cas9-NG showed lower editing efficiency than the wild-type Cas9 for the target regions ARF4T1 and ARF4T2, both followed by the NGG PAM (Figure 6B and Table S9). Mutations were induced by xCas9 in one of the three targets with the NGA PAM, ARF4T5, with the editing efficiency of 5%. Cas9-NG induced mutations in the other two NGA PAM targets, ARF4T6 and ARF4T9, with the editing efficiency of 16.7% and 3.4%, respectively (Figure 6B and Table S9).

## Discussion

In the current study, we successfully applied the natural and engineered variants of the Cas12a and Cas9 nucleases for genome editing in wheat. Even though FnCas12a was reported to be effective for genome editing in rice (Zhong et al., 2018), FnCas12a did not induce detectable mutations in our experiments, making it unsuitable for wheat genome engineering. This observation indicates that further optimization of newly CRISPR-Cas systems is usually required to improve their portability across species, even for those that belong to the same clade. On contrary, the Cas12a nuclease from *L. bacterium* was capable of inducing double strand breaks in the wheat genome, albeit at the lower rates than those previously observed in our studies for Cas9 (Wang et al., 2016; Wang et al., 2018b; Wang et al., 2019). Both the LbCas12a and engineered LbCas12a-RVR variants showed no off-target activities and ability to support highly specific genome editing in wheat.

The recovery of mutant lines with the loss-of-function mutations in the *TaGW7-B1* gene, demonstrating the expected effects on the grain size and weight traits (Wang et al., 2019), suggests that the Cas12a-based editors are effective tool for modifying the wheat genome, especially in the regions that lack the NGG PAMs recognized by Cas9. To our best knowledge, this is the first report on the usage of Cas12a for producing heritable mutations in the wheat genome with the validated effect on phenotypic traits. We also found that the efficiency of Cas12a-based multiplexed genome editing assessed in the protoplasts was not directly related to that obtained in the transgenic plants. Consistent with this observation, our prior work based on Cas9 showed that even though the protoplast-based screening is critical for selecting guide RNAs for robust genome editing, the genome editing efficiencies assessed using this approach show relatively low correlation with the number of transgenic plants carrying mutations at the target loci (Wang et al., 2018b).

We show that a crRNA array without spacers in combination with Cas12a could be applied for the multiplexed wheat genome editing. In agreement with our previous study comparing the efficiency of simplex and multiplex genome editing using CRISPR-Cas9 in wheat (Wang et al., 2018b), most Cas12a targets were edited with the efficiency equal to that achieved using the simplex guide RNAs. In addition, in the wheat protoplasts, we found increase in the GW2T6 target editing efficiency with increase in the level of multiplexing. While this trend could be explained by various factors, including the increased Cas12a recruiting capacity of the higher-level multiplexed crRNA arrays, the position of protospacers in the crRNA array or by the high accessibility of the target loci in the complex genome, it is currently hard to discern a specific reason for this observation.

Contrary to earlier work showing the improvement of genome editing efficiency after codon optimization in rice (Xing et al., 2014), the codon optimization of LbCas12a did not affect the efficiency of wheat genome editing. However, modifications introduced into the guide RNA processing system had a substantial effect on the LbCas12a’s editing ability. Even though the Cas12a nuclease is capable of processing crRNA arrays to produce mature crRNAs (Fonfara et al., 2016), its natural crRNA processing capability appears to limit the efficiency of genome editing in wheat, which could be substantially enhanced by supplementing crRNA constructs with ribozymes (Tang et al., 2017). Although most Cas12a targets had editing efficiency lower than 5%, some exhibited high mutation rates, as high as 24%, indicating that genome editing efficiency could be further improved by optimizing the CRISPR-LbCas12a system. In addition, the efficiency of Cas12a-induced genome editing could be enhanced by increasing temperature (Malzahn et al., 2019), or by improving the tolerance of LbCas12a to low temperature (Schindele and Puchta, 2020). Further studies are warranted to assess these strategies for improving the Cas12a-based genome editing efficiency in wheat.

The engineered variants of Cas12a and Cas9, which demonstrated the ability to recognize noncanonical PAMs (TATV and NG), further expand the scope of editable loci in the wheat genome. The application of engineered Cas12a variants to plant genome editing was limited (Zhong et al., 2018), and the successful modification of the wheat genome targets located next to the TATC PAM using the altered LbCas12a-RVR nuclease provides great addition to the wheat genome editing toolbox and broaden the range of species whose genomes could be modified using LbCas12a. We showed that the xCas9 and Cas9-NG nucleases targeting minimal NG PAM were effective at generating double strand breaks within the endogenous gene targets that could not be edited using other editors. Though both xCas9 and Cas9-NG recognized targets with the NG PAMs, in wheat, we observed some bias in the target preference, with Cas9-NG being more effective at targets followed by the NGH PAM than xCas9 (Zeng et al., 2019; Zhong et al., 2019).

## Conclusion

Here, we evaluated the ability of the natural (FnCas12a, LbCas12a) and engineered (LbCas12a-RVR, xCas9 and Cas9-NG) variants of the CRISPR-based DNA editors to induce mutations within the endogenous gene targets with the canonical and altered PAMs in the complex wheat genome. We demonstrated the improved target editing efficiency in the wheat genome using the LbCas12a constructs with crRNA units flanked by ribozymes. By using the LbCas12a nuclease in combination with the multiplexed RNA guides, we created stable wheat mutants with higher grain size and weight. We showed that the scope of editable loci in the wheat genome could be expanded by using the engineered LbCas12a-RVR, xCas9 and Cas9-NG nucleases recognizing targets with altered PAMs. Our study also highlights the importance of the systematic testing and optimization of newly developed CRISPR-Cas-based genome editing technologies to create a crop-specific customized toolkits to effectively implement diverse genome editing strategies for improving agronomic traits.

## Experimental procedures

### Plasmid Construction

To construct the plasmids for this study, all the DNA oligoes and fragments were synthesized by Integrated DNA Technologies (USA). All the PCR was performed using NEBNext^®^ High-Fidelity 2X PCR Master Mix (Catalog number: M0541L, New England Biolabs Inc., USA) following the manufacturer’s instructions. All DNA fragments were assembled using the NEBuilder^®^ HiFi DNA Assembly Cloning Kit (Catalog number: E5520S, New England Biolabs, USA) following the manufacturer’s instructions. All the newly constructed plasmids were confirmed by Sanger sequencing.

A plasmid with AsCas12a crRNA (CRIPSPR RNA of Cas12a from Acidaminococcus sp. BV3l6) expression cassette was firstly constructed. The NOS terminator and wheat U6 promoter in plasmid pA9Cas9sg (Wang et al., 2016; Wang et al., 2019) were amplified using primers NOS-F and DR-AsCas12aF1 (Table S1). The direct repeat sequence of AsCas12a crRNA, two BsaI cutting sites, and 7 “T” bases were added by the second round of PCR using the NOS-F and DR-AsCas12aF2 primers (Table S1). The final PCR products were subcloned into pA9FeYFP (Figure S1) between the XmaI and SacI cutting sites by replacing the eYFP and NOS terminator. Henceforth, the resulting construct is referred to as pA9Ascr.

To construct plasmid pA9LbCas12acr (9LCC for short) that expressed the humanized LbCas12a and crRNA, the plasmid pA9Ascr was amplified using the primer pair LbCrF and LbCrR. The PCR product was self-ligated using NEBuilder^®^ HiFi DNA Assembly Cloning Kit, and henceforth is referred to as pA9LbCr. The humanized LbCas12a CDS and the 3’ end NLS were amplified using the LbCas12aF and LbCas12aR primers from plasmid pY016 (pcDNA3.1-hLbCas12a). The PCR product was subcloned into pA9Lbcr to create the plasmid 9LCC (Figure S1). To add one more nuclear localization signal peptide to the 5’ end of humanized LbCas12a CDS, a primer pair SV40NLS_LbCas12a_F and LbCas12aR was used to amplify the humanized LbCas12a CDS and the 3’ end NLS. Then the resulting PCR product was amplified again using a primer pair pA9_SV40NLS_F and LbCas12aR, followed by subcloning into pA9Lbcr to create a plasmid pA9LbCas12acr-new-version, 9LCCnv for short (Figure S1). To create plasmid pA9 LbCas12aRVRcr, the plasmid pA9LbCas12acr was amplified using a primer pair pA9_SV40NLS_F and LbCas12aRVR-R, and a primer pair LbCas12aRVR-F and LbCas12aR. The resulting PCR products were ligated and amplified using a primer pair pA9_SV40NLS_F and LbCas12aR followed by subcloning into KpnI and XmaI digested pA9LbCas12acr-new-version to replace the wild type Cas12a. This created the plasmid 9LCC.

To construct pA9FnCas12acr (9FCC for short) expressing a wheat codon optimized FnCas12a and its crRNA, the FnCas12a coding sequence with NLS and 3 × HA tag on the 3’ ends was synthesized (Figure S1), and amplified using a primer pair SV40NLS_FnCas12a_F and FnCas12aR (Table S1). The resulting DNA fragment was amplified again using a primer pair pA9_SV40NLS_F and FnCas12aR to finalize the addition of SV40 NLSs. The PCR product was subcloned into pA9Ascr between KpnI and XmaI cut sites, resulting in plasmid pA9FnCas12aAscr. To change the crRNA direct repeat sequence from AsCas12a to FnCas12a, the plasmid was amplified using two primer pairs: NOS-F and FnCrR, and FnCrF and FnCas12aR. The resulting two PCR products were assembled. This created the plasmid 9FCC.

To construct the pA9LbCas12a-enhanced (9LCCe for short), the hammerhead ribozyme and hepatitis delta virus ribozyme sequences were added to the 5’ and 3’ ends of LbCas12a crRNA direct repeat, respectively, using the two rounds of PCR. Two BsaI cutting sites were embedded between the crRNA direct repeat and delta virus ribozyme. The first round of PCR was conducted using the primer pair Hammer-crF and HDVribo-crR with plasmid 9LCC as a template. The second PCR was done using the primer pair Hammer-F and HDVriboR. The PCR product was assembled with PstI +XhoI digested pMOD_B2312 (Cermak et al., 2017). The new construct, designated as pMODLBcr, was then amplified using the primer pair PvUbi1pF4 and 35SterR2, and the 9LCC was amplified using the primer pair NOS-F5 and NOS-R followed by digestion with BamHI. The first PCR product and the digested product were assembled and amplified again using primer NOS-F5 and 35SterR2. The new PCR product was assembled together with the 9LCC construct digested with XmaI and SacI to generate plasmid 9LCCe, which was then amplified using the two primer pairs, PvUbi1pF4 and PvUbi1pR1, and 35SterF2 and 35SterR2. These two PCR products were assembled and amplified using the primer pair PvUbi1pF4 and 35SterR2. The new PCR product was assembled with SpeI and SacI digested 9LCCe to form the plasmid pA9LbCas12a-enhanced2 (9LCCe2 for short). Then the plant codon-optimized LbCas12a was amplified from plasmid pYPQ230 (Tang et al., 2017) and used to replace the human codon-optimized LbCas12a in 9LCCe and 9LCCe2, henceforth pA9LbCas12a-optimized (9LCCop for short) and pA9LbCas12a-optimized2 (9LCCop2 for short), respectively.

To obtain xCas9 and Cas9-NG constructs, mutations were introduced into the maize codon-optimized zCas9 in plasmid pBUN421 (Xing et al., 2014), resulting in the pBUN421x and pBUN421NG constructs, respectively. To construct pBUN421x, zCas9 was amplified with the two primer pairs, zCas9F3 and zCas9seq4, and zCas9R3 and zCas9seq5. Both PCR products were assembled along with the synthesized DNA fragment zxCas9_741-3740. The assembled DNA fragment was amplified using the pair of primers zCas9F3 and zCas9R3, followed by assembling the resulting PCR product with XmaI and StuI digested pBUN421 to replace the wild type Cas9. To construct pBUN421NG, zCas9 was amplified using the primers zCas9F3 and zCas9seq19, and the resulting product was assembled along with the synthesized DNA fragment zCas9-NG_R. The assembled product was amplified using the primers zCas9F3 and zCas9R3, followed by assembling the PCR product with XmaI and StuI digested pBUN421 to replace the wild type Cas9.

The sequences of the wheat orthologs of six rice genes, including *OsGW2* (Song et al., 2007), *OsGS3* (Mao et al., 2010), *OsGSE5* (Duan et al., 2017), *OsAn-1* (Luo et al., 2013), *OsSPL16* (Wang et al., 2012), *OsARF4* (Hu et al., 2018b), were identified by comparing with the IWGSC RefSeq v2.0 reference genome on Ensembl Plants (https://plants.ensembl.org/index.html). The crRNA protospacers were selected from the cDNA sequences of the *TaPDS, TaGASR7* (Ling et al., 2013), *TaGW2, TaGS3, TaGSE5, TaAn-1, TaSPL16*, and *TaGW7* genes (Wang et al., 2019). Both forward and reverse sequences of the protospacers along with the 4-nucleotide 5’ overhangs were synthesized and sub-cloned into pA9FnCas12acr, pA9LbCas12acr, pA9 LbCas12aRVRcr, pBUN421x, and pBUN421NG, as previously described (Wang et al., 2018b). To construct the multiplex gene editing constructs with three, four, or eight tandem crRNA units, we synthesized one or two ultra-DNA oligonucleotides (Table S1). These oligonucleotides were assembled and amplified by PCR, and resulting amplicons were sub-cloned into plasmid 9LCCnv.

### Protoplast transformation

The wheat protoplast transient expression assay was performed as previously described with some modifications (Wang et al., 2018b). About 100 seedlings of wheat cultivar Bobwhite were grown in the dark for two weeks, shoot tissues were finely sliced and vacuumed at -600 mbar for 30 min in a 30 ml of W5 solution (0.1 % glucose, 0.08 % KCl, 0.9 % NaCl, 1.84 % CaCl_2_·2H_2_O, 2 mM MES-KOH, pH 5.7). Then the tissues were digested for 2.5 hours in a 30 ml enzymic mix containing 1.5% Cellulase R10 (from *Trichoderma viride*, 7.5 U/mg), 0.75 % Macerozyme R10 (from *Rhizopus* sp.), 0.6 M mannitol, 10 mM MES pH 5.7, 10 mM CaCl_2_ and 0.1% BSA. After digestion, protoplasts were filtered through 40 μm nylon meshes. The remaining tissues were washed with the 30 ml of W5 solution followed by filtering through the nylon meshes. The resulting cell suspension was mixed gently, and protoplasts were collected by centrifugation at 100 g for 5 min, and then washed twice with the 10 ml of W5 solution. The final protoplast pellet was re-suspended in the 5 ml of W5 solution, cell count was estimated using a hemocytometer. The re-suspended protoplasts were kept on ice for 30 min to allow for the natural sedimentation. Then protoplast cell count was adjusted to 10^6^ cells/ml in MMG solution (0.4 M mannitol, 15 mM MgCl_2_, 4 mM MES, pH 5.7).

The 10 µg of plasmid DNA and 100 µl of protoplasts were mixed with the 130 µl of PEG solution (40% (W/V) PEG 4000, 0.2 M mannitol and 0.1M CaCl_2_). After 30-min incubation at room temperature in the dark, 500 µl of W5 solution was added. The protoplasts were collected by centrifugation at 100 g for 2 min, re-suspended in 1 ml of W5 solution and incubated in the dark at room temperature. The transformation efficiency was assessed by counting the fraction of fluorescent-positive protoplasts transformed with pA9mRFP. Protoplasts were collected 48 h after transformation and DNA was isolated with PureLink Genomic DNA Mini Kit (Thermo Fisher Scientific, Catalog number: K182002) following the manufacture’s protocol.

### Gene editing efficiency calculation by the next generation sequencing (NGS) of PCR amplicon library

To detect mutations induced by the CRISPR-Cas12a or CRISPR-Cas9 variants, genomic regions harboring the crRNA targets were amplified by PCR. The Illumina’s TruSeq adaptors on both ends of the amplicons were added using two rounds of PCR as described (Wang et al., 2016). PCR products were purified with MinElute PCR Purification Kit (Qiagen), pooled in equimolar ratio, and sequenced on a MiSeq Sequencer using the MiSeq Reagent Nano Kit v2 (500 cycles, 2 x 250 bp run) at the K-State Integrated Genomics Facility. The Illumina reads passing quality control were aligned to the wild-type reference sequences of targeted genes. The gene editing efficiency was calculated by dividing the number of mutated reads to the total number of all aligned reads.

### Transgenic plants regeneration and genotyping

Wheat immature embryo transformation and plant regeneration were performed as previously described (Saintenac et al., 2013). To isolate DNA, leaf tissues were sampled and homogenized in 500 μL of TPS buffer, then incubated for 20 min at 75 °C. After centrifugation for 5 min, 140 μL of the supernatant was mixed with 140 μL isopropanol and incubated for 20 min at room temperature. DNA was precipitated, washed with 70% ethanol, and re-suspended in 100 μL of deionized water.

The presence of CRISPR/Cas12a or CRISPR-Cas9 variants constructs in the transgenic plants was validated by PCR using four pairs of primers amplifying different regions of the Cas12a and crRNA expression cassettes (Table S1). The CRISPR/Cas12a-induced mutations were examined only in the plants showing the presence of three PCR products. The mutations were detected using the NGS-based procedure described above.

### Plant growth and grain morphometric data collection

The CRISPR-Cas12a induced mutant plants were grown and phenotyped as described previously with slight modifications (Wang et al., 2018c). Briefly, the T1 generation plants were grown in a growth chamber under 16-h light / 8-h dark cycle. The temperature was set to 24 °C during the day and 20 °C during the night. The T_2_ generation plants were grown in the Kansas State University’s greenhouses under natural conditions supplemented by additional light sources to maintain 16-h light / 8-h dark cycle. The room temperature was set as 24 °C during the day and 20 °C during the night. Three main spikes from each plant were harvested separately. A MARVIN seed analyzer (GTA Sensorik GmbH, Germany) was used to measure the grain morphometric traits (grain width, length, area), and thousand grain weight for each spike. The mean of three spikes from each plant was calculated and used for further analyses.

### Statistical analysis

The two-tailed Student’s *t*-test was applied to assess the significance of differences between the simplex and multiplex gene editing efficiencies in the wheat protoplasts.

## Author Contributions

W.W. conducted the gene editing experiments in protoplasts and the genotyping and phenotyping of transgenic plants, generated and analyzed the next-generation sequencing and phenotyping data, drafted the manuscript; B.T. conducted plant transformation experiments, provided the GFP transgenic wheat, helped the design of gene editing experiments; Q.P. analyzed gene editing events using next-generation sequencing; Y.C. performed biolistic transformation of wheat embryos; F.H. developed pipeline and helped for NGS data analyses; A.A. designed experiments for NGS analysis of editing events and performed NGS; H.T. performed biolistic transformation of wheat embryos with the gene editing constructs and coordinated project; GB provided Sanger sequencing; and E.A. conceived idea, coordinated project, and wrote the manuscript. All authors revised the manuscript and approved final version of the manuscript.

## Acknowledgements

This project was supported by the Agriculture and Food Research Initiative Competitive Grants 2017-67007-25932 and 2020-67013-30906 from the USDA National Institute of Food and Agriculture, and grants from the Bill and Melinda Gates Foundation and Kansas Wheat Commission. We thank Dwight Davidson for the greenhouse management and phenotyping data collection. We thank Dr. Feng Zhang, Dr. Yiping Qi, Dr. Qi-Jun Chen, and Dr. Daniel Voytas for providing the plasmid pY016, pYPQ230, pBUN421, and pMOD_B2312, respectively.

## Conflict of interest

The authors declared that they do not have conflict of interests.

## Supporting Information

Figure S1. The sequences of plasmids applied in this study.

Figure S2. Gene editing efficiency comparison of single target and multiple target CRISPR-LbCas12a constructs.

Figure S3. The off-target blast hits of 12 LbCas12a targets against Chinese Spring RefSeqv1.0.

Table S1. The primers and DNA oligoes used in this study.

Table S2. The NGS-based gene editing efficiency analyses of CRISPR-LbCas12a constructs in wheat protoplasts.

Table S3. The NGS-based gene editing efficiency analyses of CRISPR-FnCas12a constructs in wheat protoplasts.

Table S4. The NGS-based gene editing efficiency analyses of CRISPR-Cas9 constructs in wheat protoplasts.

Table S5. The NGS-based gene editing efficiency analyses LbCas12a multiplex constructs in wheat protoplasts.

Table S6. The NGS-based gene editing efficiency analyses of improved LbCas12a constructs in wheat protoplasts.

Table S7. The NGS-based gene editing efficiency analyses of LbCas12a-RVR constructs in wheat protoplasts.

Table S8. The NGS-based off-target analyses of LbCas12a and LbCas12a-RVR constructs. Table S9. The NGS-based gene editing efficiency analyses of Cas9 variants in wheat protoplasts.

## Notes

### Competing Interest Statement

The authors have declared no competing interest.

